# Population-scale prediabetic assessment using HbA1c from menstrual blood

**DOI:** 10.1101/758805

**Authors:** Lakshminarayan Srinivasan, Reshma Khilnani, Philip Fung, Melissa Low, Urano Esquivel, Bridget White, Gurbir Sidhu, Jesus Rangel, Shion A. Lim, Leanna S. Sudhof, José O. Alemán

## Abstract

CDC-recommended diabetes prevention programs aim to detect and reverse disease in the one-third of Americans with prediabetes, but high-compliance serial assessment of percent hemoglobin A1c (HbA1c) remains a barrier to delivering this vision at population scale. Venous phlebotomy is challenging for busy or resource-constrained patients.

In this paper, we introduce the first-ever quantitative diagnostic test based on menstrual fluid, which allows HbA1c quantification from self-collected mailed tampons.We demonstrate that menstrual HbA1c is comparable to venous HbA1c in the diagnosis of prediabetes with the standard threshold of 5.7. We also demonstrate accuracy, precision, stability, and interference testing. Finally, surveying subjects reveals strong preference for menstrual HbA1c in quarterly testing. These findings suggest that menstrual HbA1c can be a key tool in addressing prediabetes at population scale.

## 1 INTRODUCTION

The impact of diabetes at the population level is immense. In the United States (US) alone, 9.4% of Americans have diabetes ^[1]^, and an additional 33.9% of people have prediabetes ^[2]^, distributed evenly across adult age groups. An estimated 23.8% of patients with diabetes and 88.4% of patients with prediabetes are entirely unaware of their diagnosis ^[1] [2]^. The cost of caring for patients with diabetes accounts for 14% of overall healthcare expenditure in the US, even without accounting for the elevated risks of heart attack and stroke attributable to diabetes^[3]^. At the same time, the cost of insulin has skyrocketed, reducing access to adequate treatment for diabetes. As risk factors for diabetes include ethnicity, age, obesity, and diet, the prevalence of prediabetes varies geographically, exceeding 50% in some states (Figure 1A).

**FIGURE 1.**
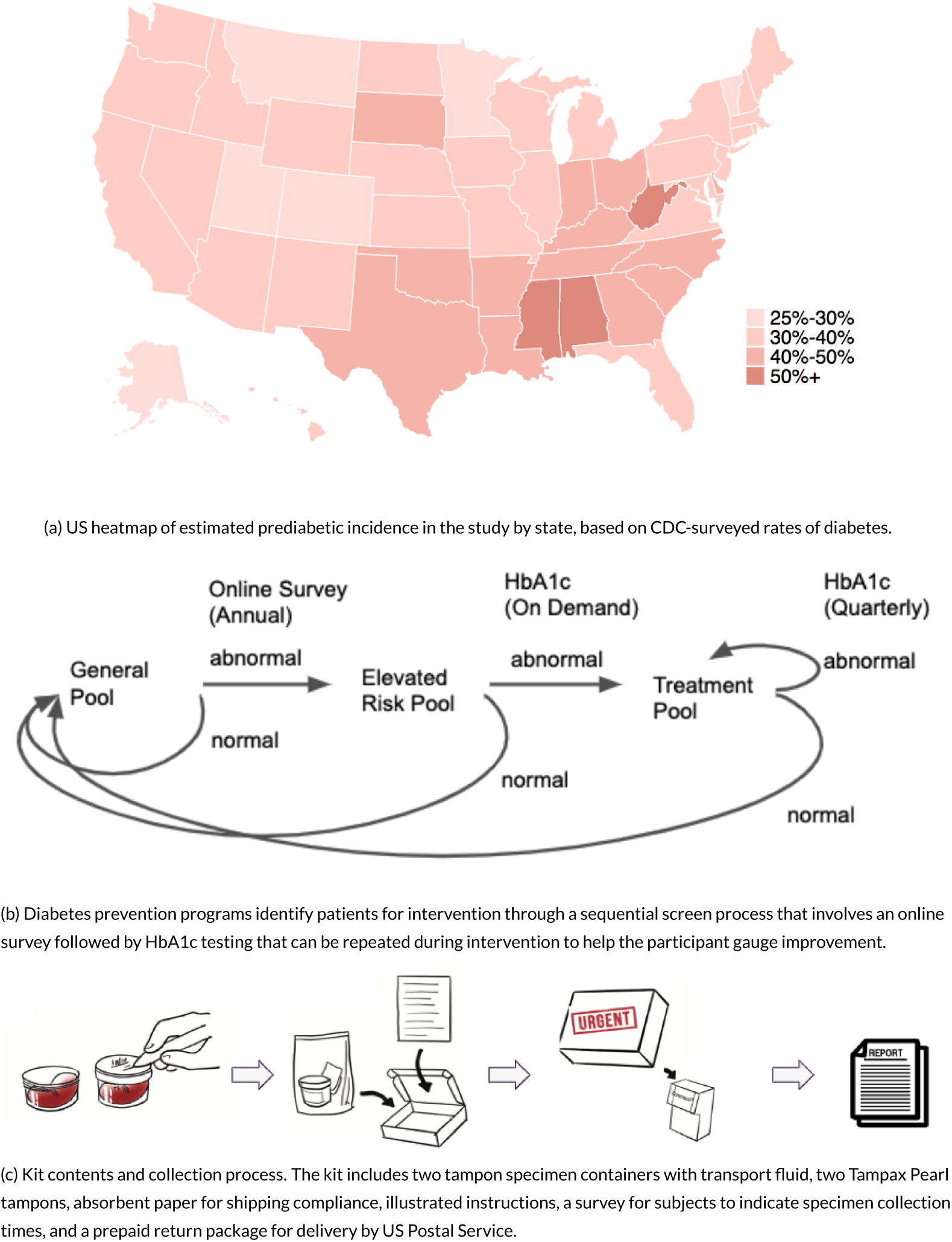
Diabetes prevention program based on menstrual fluid.

Led by the CDC Diabetes Prevention Program, the American Diabetes Association ^[4]^, and other government initiatives, there is growing recognition that high-risk patients can be guided to avoid diabetes entirely ^[5] [6]^, by screening the general population, and directing intensive lifestyle intervention to patients at elevated risk for diabetes (Figure 2). In the CDC prevention program, the general population is first screened for elevated diabetes risk with a brief online survey that searches for risk factors including age, sex, history of gestational diabetes, first-degree relative with diabetes, elevated blood pressure, physical activity, weight and height^[7] [8]^. Resources are then directed in a periodic fashion at patients within the pool that demonstrate elevated percentage hemoglobin A1c (HbA1c), a routine blood test of a glycosylated hemoglobin isoform ^[9,10,11]^ which is currently the gold standard for diagnosing and monitoring prediabetes (HbA1c 5.7-6.4) and diabetes (HbA1c > 6.4) ^[12]^. Regular HbA1c testing can help determine whether behavioral intervention is working at the patient and population levels, and also guides physician reimbursement under the current Quality Payment Program (CMS ID 122v5).

**FIGURE 2.**
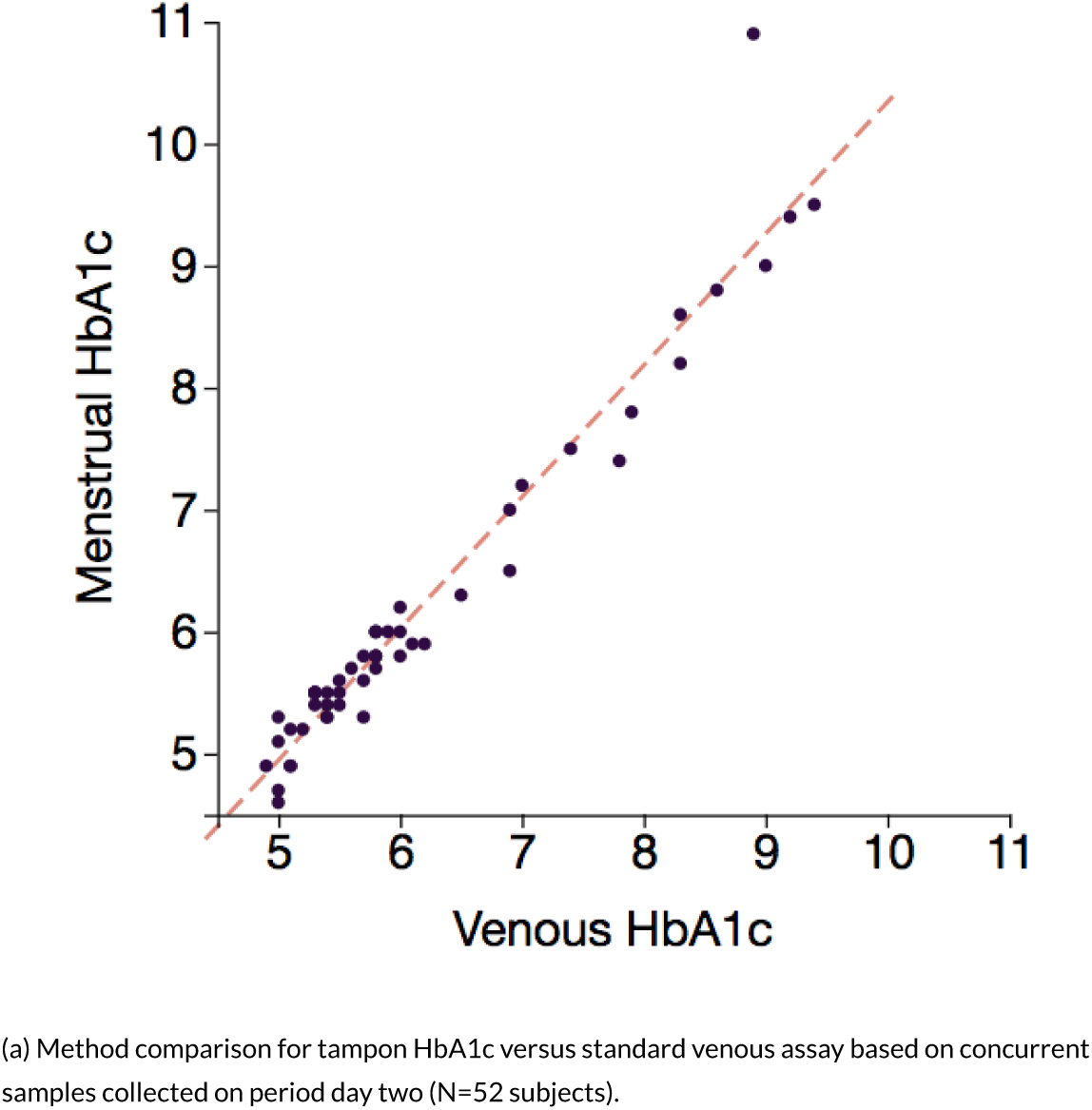
Menstrual HbA1c versus traditional venous testing.

HbA1c is assessed by venous phlebotomy, collected into an EDTA-coated container, and coupled with an assay. Some assay categories include enzymatic reaction, antibody-latex agglutination, capillary electrophoresis, and high-performance liquid chromatography (HPLC). Because venous phlebotomy traditionally requires advancing a needle into the antecubital vein in sterile fashion, it is typically performed by licensed professionals at centralized phlebotomy sites within hospitals or outpatient centers.

Compliance with venous phlebotomy orders is poor among patients with diabetes. Among 27,096 patients aged 19-50 with known diabetes at Kaiser Permanente Northern California, 21.5% failed to arrive for venous HbA1c testing within 30 days of receiving an order^[13]^. The compliance rate decreased further without the benefit of a face-to-face visit, and persisted regardless of race, socioeconomic status, or English fluency. The contributors for low phlebotomy compliance are multifactorial. Phlebotomy requires the patient to take time off from work, and the economic impact of lost wages at population scale could be substantial. Needle phobia ranges from 10% to 70% prevalence depending on specific context ^[14]^. This phobia is exacerbated by a history of traumatic exposure to needles, which would make compliance with quarterly venipuncture even more challenging. Venous access is also harder to establish in diabetic patients and patients with a variety of related risk factors include obesity, chronic illness, vasculopathy, edema, and multiple prior hospitalizations ^[15]^.

Capillary collection with a self-administered fingerstick is an over-the-mail alternative to venous phlebotomy for assessment of HbA1c. Capillary blood can be collected as dried blood spots on Whatman paper ^[16]^, into aqueous transport buffer^[17]^, or tested at home with point-of-care devices^[18]^. Both Whatman and aqueous transport options are robust to temperature excursions during shipping. While capillary collection has been used extensively in public health research, it has not yet been translated into large-scale screening for diabetes wellness programs in the US. One contributing factor may be the need for otherwise healthy patients to periodically and repeatedly overcome the pain and fear associated with self-collecting fingerstick blood. The general and prediabetic population involved with a lancet-based screening effort would not be as familiar with lancets as patients with diabetes who self-collect fingerstick blood multiple times per week. Another limiting factor may be the need for laboratories to carry specialized equipment and assay calibration for processing either dried blood spots or capillary transport tubes.

Menstrual blood is a new specimen type that could overcome some limitations of phlebotomy and capillary collection in population-scale testing. It is accessible to the large prediabetic population of women age 18-50, and prediabetes prevalence is uniformly distributed across this age range. Most premenopausal women menstruate multiple times per year. Women are also practiced in self-collecting menstrual blood onto cotton-based tampons or sanitary pads which are both regulated FDA Class II devices ^[19]^. Tempering these potential advantages, some conditions in premenopausal women will preclude use of menstrual blood, including prior hysterectomy, hormonal contraception, and other causes of oligo-or amenorrhea.

The composition of menstrual fluid includes systemic blood, endometrial, cervical and vaginal epithelium; and various bacteria and Candida species. Proteomic analysis demonstrates substantial overlap with venous blood, although menstrual fluid also contains proteins not seen in venous blood or vaginal fluid ^[20]^. Endogenous fibrinolytics including plasmin in menstrual blood inherently anticoagulate the blood constituents, which facilitates collection and transport ^[21]^.

In this paper, we demonstrate the feasibility of determining HbA1c at population scale using menstrual blood collected by tampon via mail (Figure 1C). First, we demonstrate equivalence between menstrual and venous HbA1c in the diagnosis of prediabetes. Second, we characterize the time delays and temperature conditions expected from over-the-mail tampon collection in the US. Third, we demonstrate sample stability under representative delay and temperature conditions. Fourth, we show that menstrual HbA1c is robust to common vaginal interferents. Finally, we survey participants to demonstrate an overwhelming preference for this over-the-mail menstrual assay versus conventional venous phlebotomy.

### 1.1 Results

#### 1.1.1 Method Comparison

HbA1c values were concordant between venous blood and menstrual blood (Figure 2A). Each point in this graph represents the most recently collected tampon in a kit together with the corresponding venous value. Linear regression demonstrated a slope of 1.08 (1.00, 1.15) with y-intercept of -0.45 (−0.91, 0.01) and R2 value of 0.95. A histogram of menstrual HbA1c errors across all tampons within kits regardless of tampon ordering (Figure 2B) aggregates errors computed versus both venous and capillary HbA1c. This graph demonstrates that only 1 out of 52 subjects (less than 3% of subjects) exhibited errors that exceeded 10% relative to concurrently drawn venous blood.

#### 1.1.2 Outlier Analysis

Errors that exceeded 10% relative to concurrently drawn venous blood were seen in only 1 out of the 52 subjects. This single error demonstrated tampon HbA1c of 10.5 versus venous HbA1c of 8.9. Because the true HbA1c value here was in the diabetic range, this positive error in tampon HbA1c did not contribute to error rates for prediabetic or diabetic screening. We also examined a second tampon in this kit, and 8 preceding tampons over a 6 month period preceding this kit from the same subject with various transport buffers. The second tampon from this kit also demonstrated an erroneously high value of 10.7. However, all 8 preceding tampons from previous months for this subject demonstrated errors less than 10% relative to concurrently drawn venous values, in the range of the most recent venous value included in the analysis (8.9). This suggests a correlated source of error, although the source of this error remains speculative. Temperature was normal, as temperature logs on this kit demonstrate typical temperature in the range 8-25C. High triglycerides may be the candidate explanation, as the most recent venous sample demonstrated an extremely high venous triglyceride of 1420 mg/dL (measured using a 1:2 dilution to bring value within range of the CardioCheck point-of-care device). While triglycerides are a known interferent, further work is needed to establish a definitive source of error. Because tampon HbA1c errors are rare, further exploration of error characteristics would require large population sizes on the order of thousands of patients.

#### 1.1.3 Inter-Tampon Consistency

HbA1c demonstrated consistency across tampons within the same kit (Figure 2C). The same tampons from method comparison (Figure 2A) were plotted against the second-most recent tampon. Linear regression demonstrated a slope of 0.96 (0.91,1.00) with y-intercept of 0.25 (−0.04, 0.54) and R2 value of 0.98.

As an alternate representation of consistency (Figure 2D), the unsigned difference in HbA1c between the two tampons from each kit was computed, and a distribution was formed across all kits. Data in this graph are not binned; HbA1c values are measured to one significant digit. The distribution of these differences shows that 95% of kits demonstrated an inter-tampon discrepancy of 0.5 or less, and the maximum discrepancy measured was 0.6.

**FIGURE 2.**
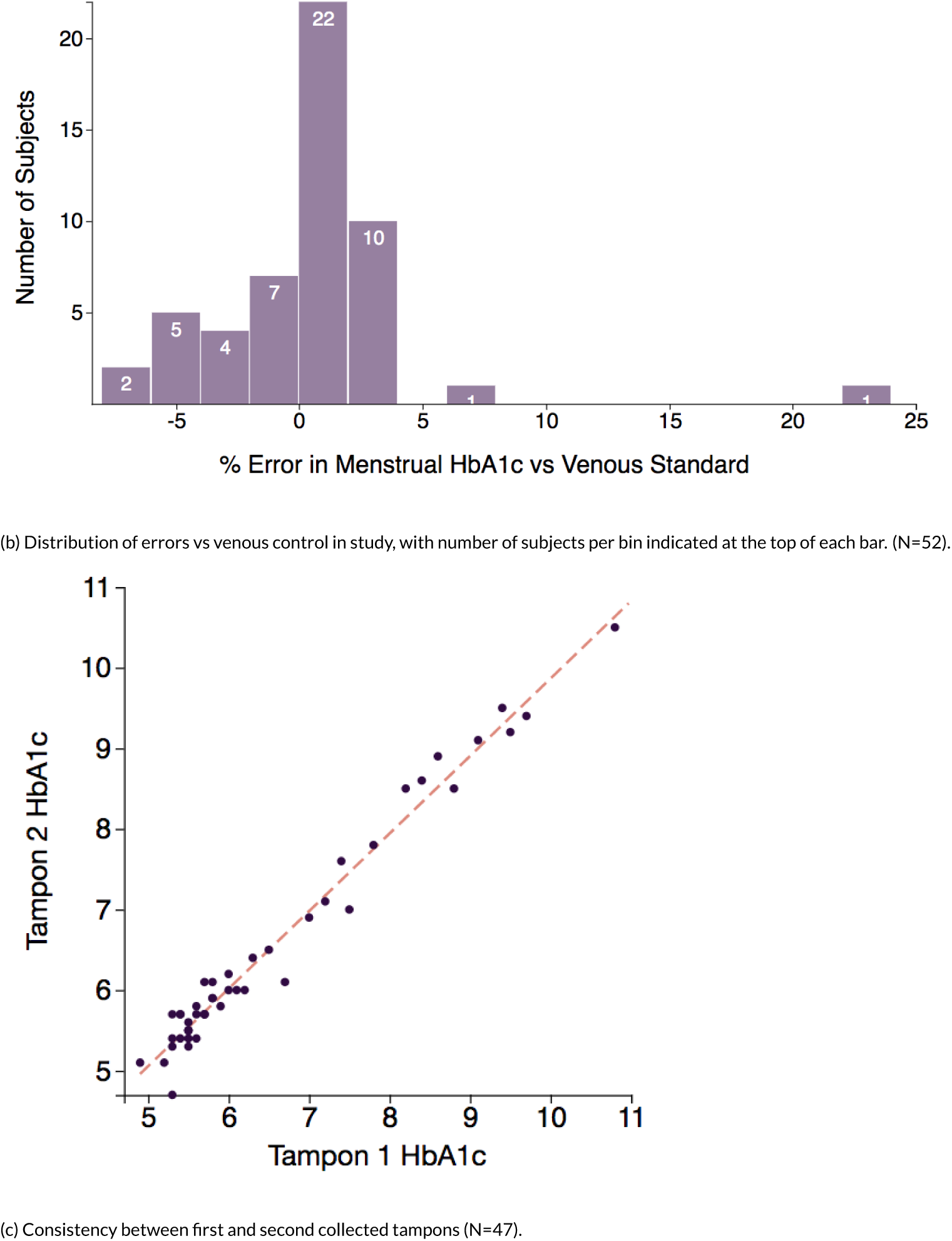
Menstrual HbA1c versus traditional venous testing.

**FIGURE 2.**
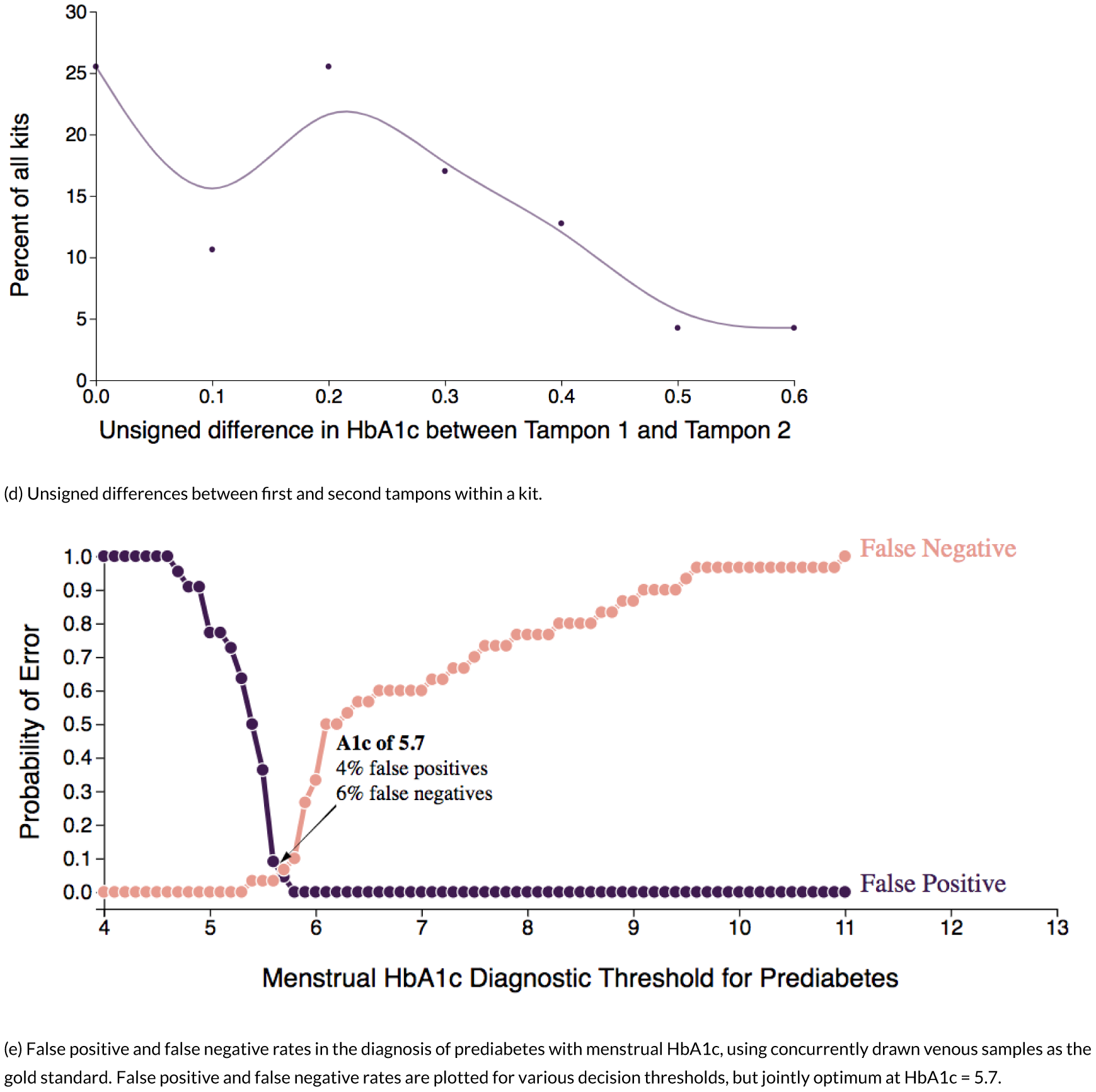
Menstrual HbA1c versus traditional venous testing.

#### 1.1.4 Diagnostic Accuracy

To evaluate whether menstrual blood performs comparably to venous blood in the diagnosis of prediabetes (Figure 2E), we compared false positive and false negative rates across a variety of proposed diagnostic thresholds for menstrual HbA1c. The diagnostic threshold that minimized the sum of both errors was 5.7, identical to the standard venous HbA1c diagnostic threshold for prediabetes. False positive and false negative rates were jointly optimum at the decision threshold of 5.7, measuring 4% and 6% respectively.

#### 1.1.5 Stability and Interference

To determine realistic stability testing conditions, an assessment of expected delay times and temperatures was performed by mailing 123 kits to subjects in 30 states. The geographic distribution of subject participation (Figure 3A) in our logistics study coincidentally approximates the intensity of prediabetes by state. An analysis of non-negotiated non-volume-discounted shipping costs for commercial services that ship within five days (Figure 3B) demonstrates that USPS priority mail typically achieves an optimum price-to-shipping ratio, with three-day shipping at pricing around $10 per kit. USPS priority mail was employed for subsequent logistics experiments.

**FIGURE 3.**
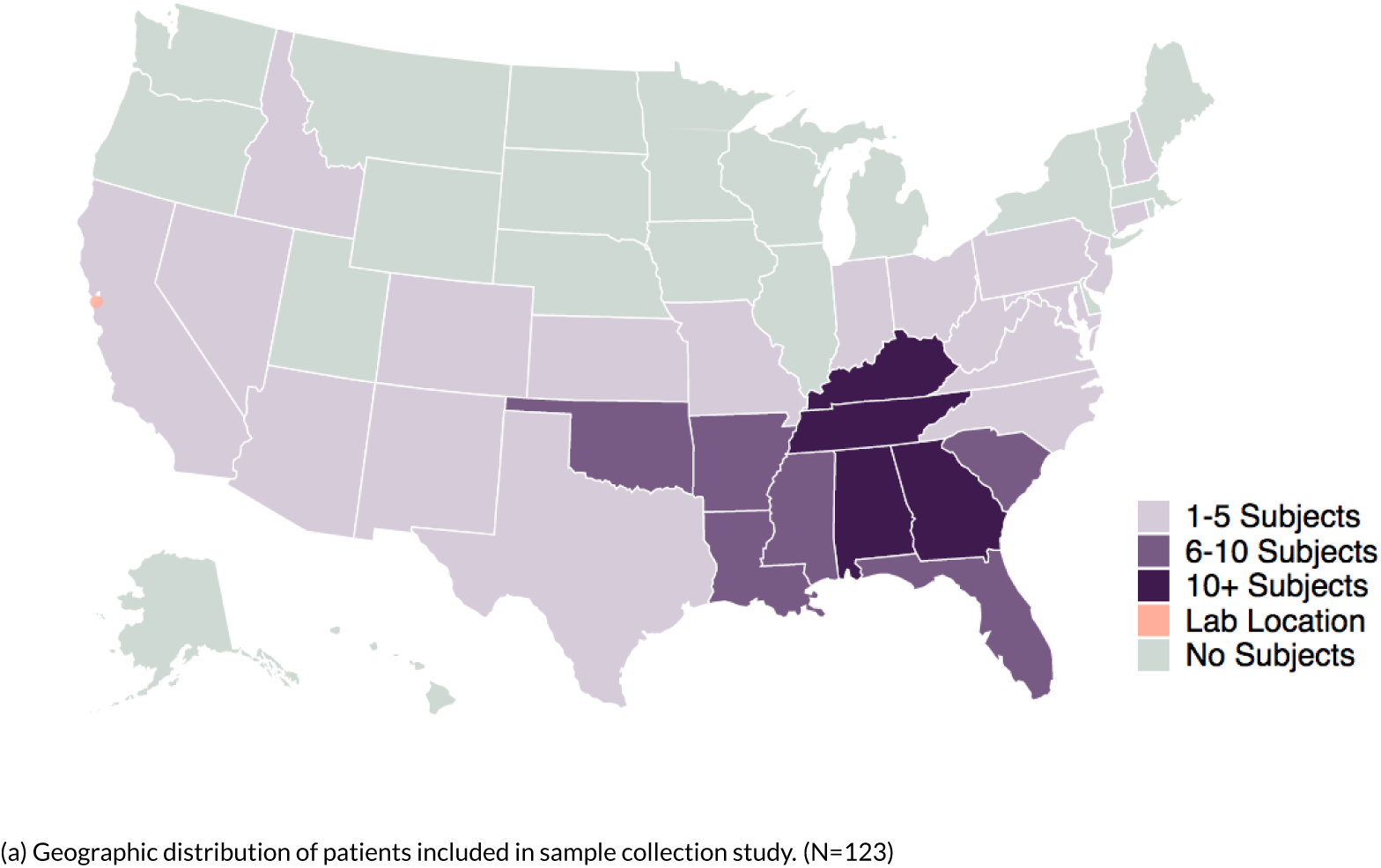
Menstrual HbA1c versus traditional venous testing.

Aggregate statistics on the duration spent in each phase of collection (Figure 3C) demonstrates that the latency between collecting the tampon and shipping it (purple) contributes to more than one-third of the specimen age (row labeled AVG), requiring nearly 2 days on average. Shipping times were typically between 1-3 days. Laboratory processing ranged from less than 1 day to as many as 5 days in the case of long weekends; this latency could be substantially reduced at scale by scheduling clinical laboratory coverage. Digital temperature data (Figure 3D) demonstrated that unrefrigerated shipping via USPS predominantly occurs at 8-25 °C with less than 15 hours spent at temperatures between 25-31 °C, and less than 6 hours spent at temperatures of 31-37 °C.

**FIGURE 3.**
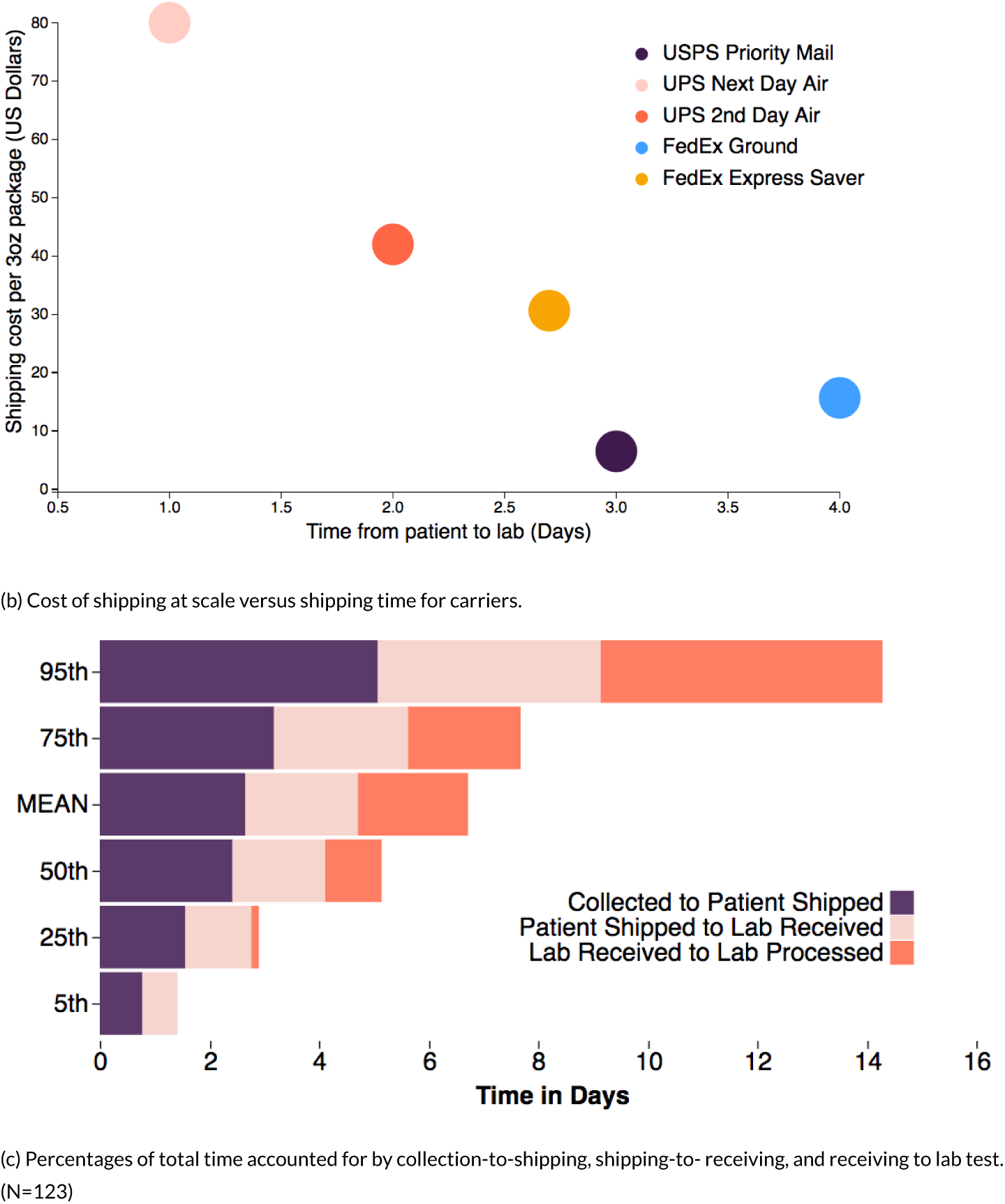
Menstrual HbA1c versus traditional venous testing.

**FIGURE 3.**
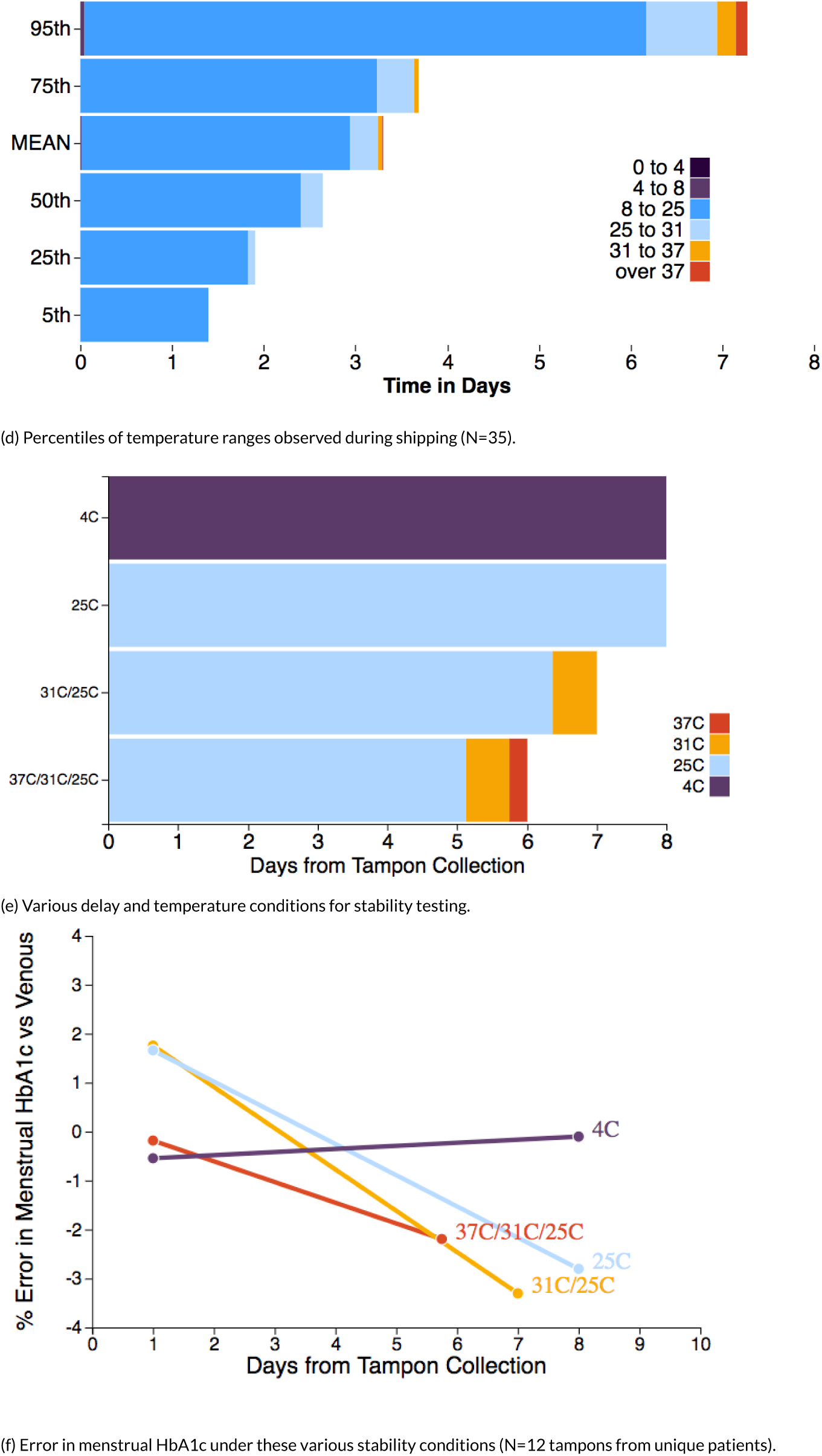
Menstrual HbA1c versus traditional venous testing.

**FIGURE 3.**
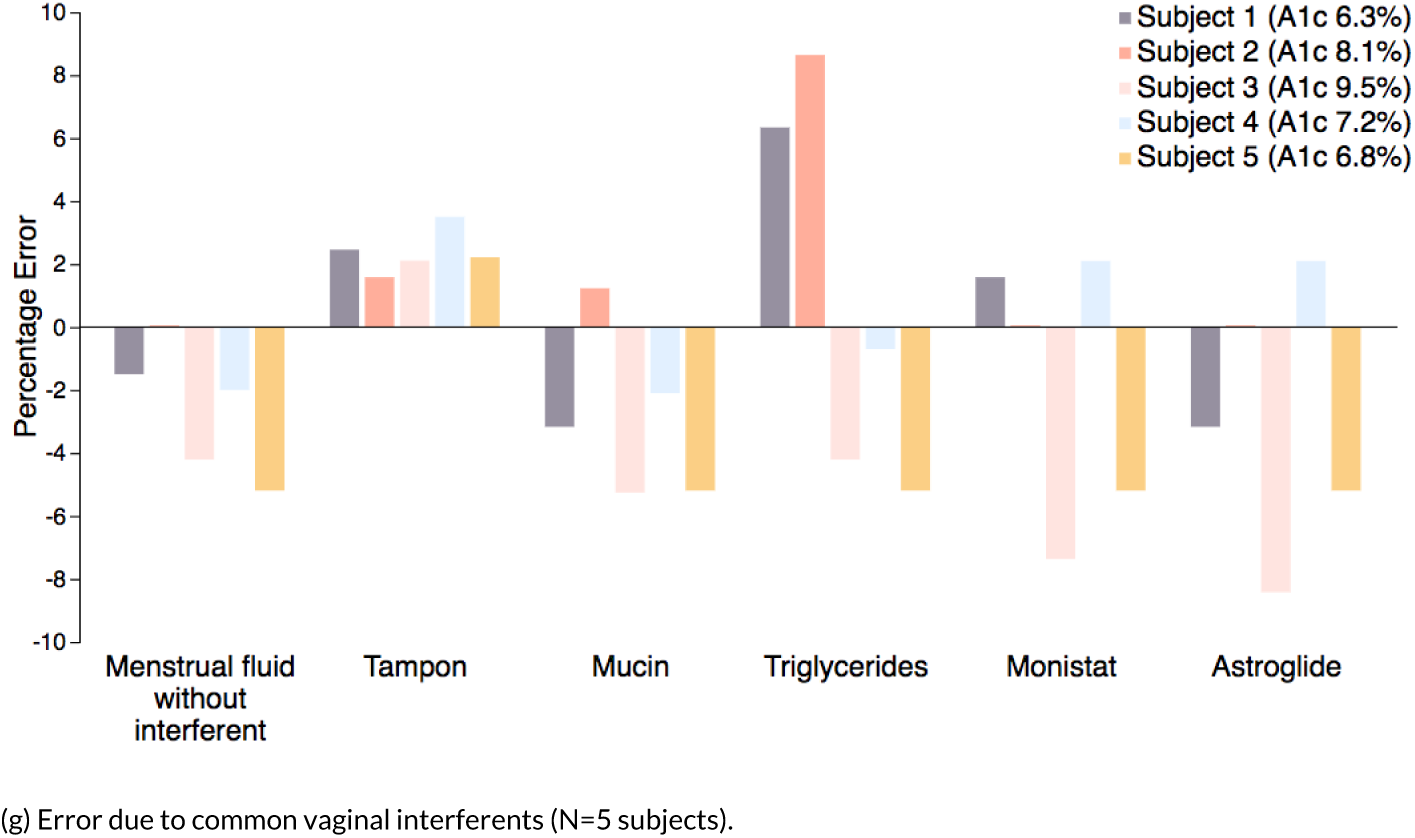
Menstrual HbA1c versus traditional venous testing.

Based on this shipping data, four simulated stability challenges were prepared (Figure 3E), including 8 days at 4 °C (first row), 7 days at 25 °C (second row), 7 days including 15 hours at 31 °C close to time of sample receipt and otherwise 25 °C (third row), and 6 days including 6 hours at 37 °C, 15 hours at 31 °C, and 5 days 3 hours at 25 °C (fourth row). For all conditions, there was a variable period of time less than 18 hours from collection to sample receipt during which the temperature sensor did not exceed 25 °C, and this transit time was included in the total time listed. Across the variety of simulated delays and temperatures, errors in HbA1c were contained within +/- 4% relative to venous HbA1c (Figure 3F).

Based on this shipping data, four simulated stability challenges were prepared ranging from 4 °C through 37 °C (Figure 3E). Across the variety of simulated delays and temperatures, errors in HbA1c were contained within +/- 4% relative to venous HbA1c (Figure 3F).

We also examined the effect of various interferents on menstrual blood collected by tampon. We exogenously spiked these interferents into samples from 5 unique subjects, each of whom provided tampons and EDTA venous blood (Supplementary Figure 1). For HbA1c assessed on a tampon-extracted menstrual sample without interferents, errors were negative and within 6%. Next, EDTA-treated venous blood applied to fresh tampon fragments demonstrated a small but systematically positive error in HbA1c resulting from interaction between the venous samples and tampon, all within 4% error. The remaining interferents demonstrated total errors within +/-10%. The rank ordering of error severity between subjects was generally preserved across interferents.

#### 1.1.6 Patient Preference

To understand patient preference for menstrual testing versus conventional venous testing, we surveyed 63 normal and prediabetic subjects based on venous or capillary HbA1c less than 6.5 (Figure 4). A total of 81 subjects were invited to participate, and responses from the 63 subjects were collected within an 8-day period.

**FIGURE 4.**
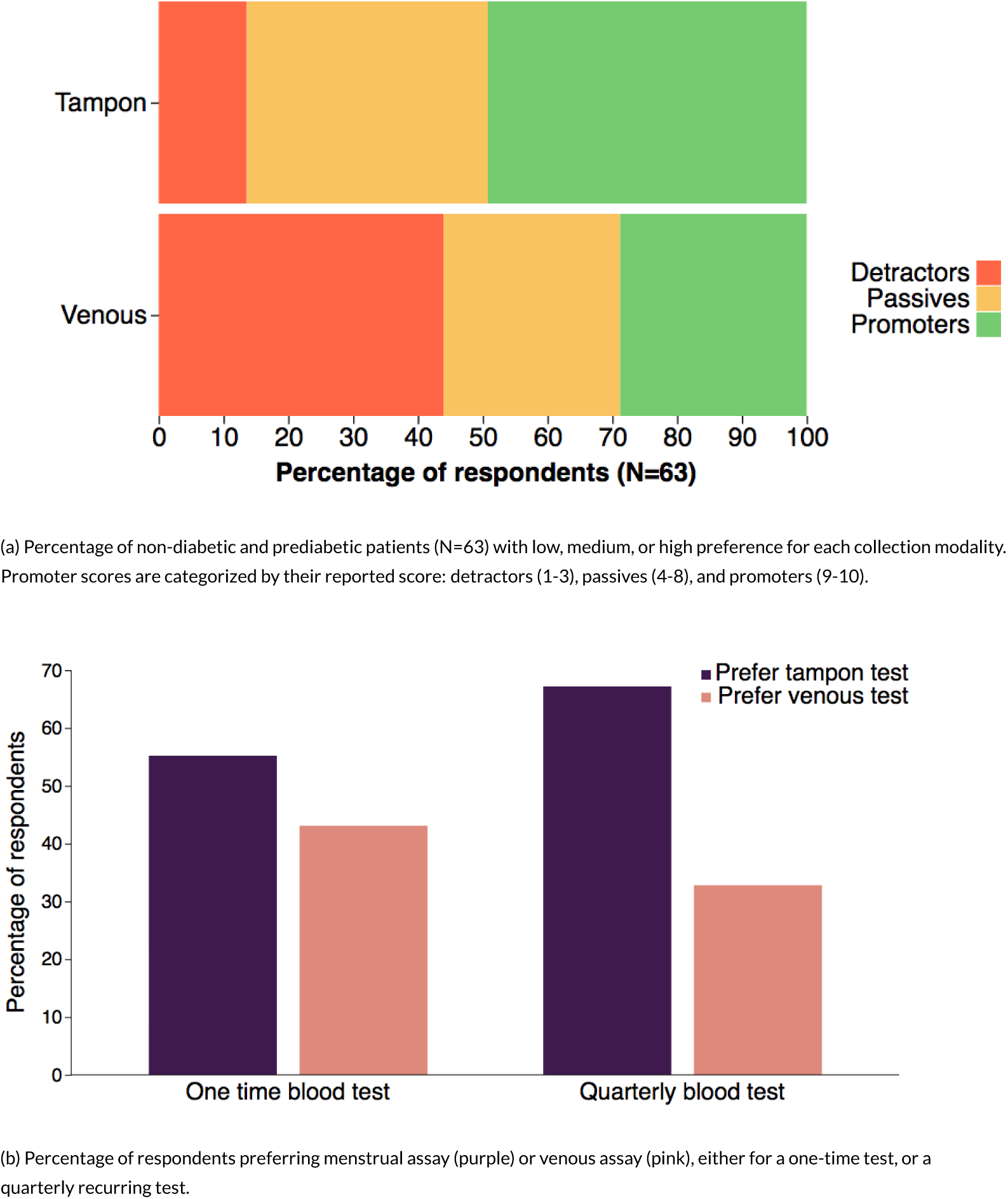
Menstrual HbA1c versus traditional venous testing.

We first asked subjects to indicate how likely they were to recommend menstrual and venous testing (Figure 4A). Preference was rated on a scale from 1 to 10, where 10 indicates very likely to recommend, and then bucketed into detractors (1-6), passives (7-9), and promoters (9-10). For venous testing, there were 26 detractors, 18 passives, and 19 promoters. For menstrual testing, there were 10 detractors, 22 passives, and 31 promoters. Differences in frequency of each preference category were compared between menstrual and tampon options using a two-proportion z-test. The percentage of detractors were significantly less for menstrual versus venous testing (p=0.0019). The percentage of promoters was significantly larger (p=0.015) for menstrual versus venous testing. The difference in percentage of passives was not statistically significant (p=0.23).

We then examined which method was more conducive to recurrent testing, using a two-choice forced selection between menstrual and venous collection (Figure 4B). Subjects were first asked to indicate their selection for a one-off test, and next to indicate their selection for a quarterly recurring test. For the one-off test, 36 subjects preferred tampon and 26 subjects preferred venous. For the quarterly test, 42 subjects preferred tampon and 20 subjects preferred venous. In both cases, menstrual collection was preferred over phlebotomy, and this preference for menstrual collection was statistically significant in the setting of quarterly testing based on one-proportion z test (p=0.0081) and did not meet significance with one-off testing (p=.26).

## 2 DISCUSSION

Diabetes prevention programs can avert the growing crisis of diabetes. The Prediabetes Risk Test by the National Diabetes Prevention Program provides a first step for patients to self-assess their risk of type 2 diabetes[4]. However, these programs also require a convenient method to screen and monitor HbA1c at population scale in patients who may be otherwise healthy and thus may not access healthcare regularly. In this paper, we introduced the first reported menstrual blood diagnostic test, and validated its use in quantifying HbA1c through tampons collected by patients and shipped at room temperature via priority mail. We also showed that our assay performs similarly to venous blood in the diagnosis of prediabetes using the standard 5.7 HbA1c diagnosis threshold for prediabetes. Subjects demonstrated a preference for a menstrual diagnostic over standard venous phlebotomy when quarterly testing was necessitated. This method meets a clinical need for the general and prediabetic population who would not be as comfortable with lancet-based tests as patients with diabetes who self-collect fingerstick blood multiple times per week^[22]^.

Remarkably, there is little prior research characterizing the material and chemical properties of menstrual blood. Some studies use transcriptomic or spectrometric techniques to distinguish menstrual and venous blood in the context of forensic work ^[23,24]^. With the rise of new high-throughput assays of various kinds, we anticipate a growing interest in the diagnostic utility of menstrual blood. Fortunately, prior exploration in hematology and clinical chemistry provide an extensive roadmap for the systematic analysis of menstrual blood.

While our work represents a first-in-woman demonstration of a menstrual blood diagnostic assay, further work is needed to expand this technology across menstrual sanitary products. The material composition of modern pads and tampons are varied and largely undocumented^[25]^, and special consideration to assay compatibility may be needed with each new tampon or pad assay. Because preference for pads and tampons varies, development of a pad-based assay would substantially widen the impact of menstrual assays.

The novelty and practicality of sample collection via tampons will likely resonate with patients who will for the first time be able to leverage a traditionally stigmatized and bothersome bodily fluid in achieving better health. Furthermore, improved screening of reproductive-aged women has the potential to optimize pre-conception health and thus decrease the adverse pregnancy outcomes associated with pregestational diabetes, such as congenital malformations, preterm birth, cesarean delivery, and perinatal morbidity and mortality ^[26]^. Increasing diabetes screening rates in women of reproductive age could also positively impact the health of their families, since maternal type 2 diabetes is associated with an increased risk of type 2 diabetes in offspring ^[27]^, and since women as caregivers may transmit lifestyle changes, particularly better nutrition, to their children ^[28]^.

While the menstrual HbA1c assay provides a compelling scalable solution for population-scale diabetes prevention programs, the potential for menstrual assays is wider. Our hope is that our work on this menstrual assay will enable further scientific understanding of menstrual blood and its use as a less invasive screening modality in menstruating individuals everywhere.

## 3 METHODS

### 3.1 Subject recruitment

Studies were approved under Western Institutional Review Board (WIRB) Protocol Number 20172117. Subjects participated remotely via mail or locally using mobile phlebotomists. All subjects were recruited via online advertisements placed through Google Ads, Facebook Ads, and Craigslist. Patients received SMS and email reminders starting the day before their expected period as described on their registration form. Both illustrated text and video were used for collection instructions. Subjects were compensated via Amazon gift card, $30 for remote donors who provided fingerstick capillary blood, and $100 for local donors who underwent more invasive venous phlebotomy.

Consent was performed through an online enrollment form, combined with a phone call and digital signature. Once donors were consented, they were able to donate multiple times.

### 3.2 Sample Procurement and Processing

#### 3.2.1 Capillary Blood Self-Collection

Subjects self-collected capillary blood on the second day of the menstrual period, concurrent with tampon collection. Capillary collection was performed by fingerstick with a lancet (Unistick 3, Owen Mumford, 30G, 1.5mm) using the Biorad variant II capillary collection device (Bio-Rad catalog number 196-2050, BioRad 510(k) K142448), including two plastic capillaries (5 uL each) and sample preparation vial. The vial contains 1.5 mL of hemoglobin capillary capture system (HCCS) reagent, an aqueous solution of EDTA and potassium cyanide.

After washing their preferred hand in warm water for one minute, subjects were instructed to clean their index finger with an alcohol swab, and lancet eccentrically into the soft tissue overlying the distal phalanx. By massaging from palm to finger, subjects were instructed to express enough blood to fill two 5 uL plastic capillaries before placing them into the sample preparation vial and shaking vigorously until the capillaries were fully rinsed.

Capillary HbA1c values were not used for method comparison. Instead, they were used to select prediabetic and normal patients for the experience survey.

#### 3.2.2 Venous Blood Draw

Mobile phlebotomists were dispatched to local donors for venous blood draw on the second day of the menstrual period, concurrent with tampon collection. A minimum of 4 mL of venous blood was drawn using standard purple-top EDTA vacutainer tubes. Venous blood was transported by mobile phlebotomist at ambient temperature to the lab within 6 hours of blood draw.

#### 3.2.3 Tampon Collection

Subjects collected 2-3 consecutive tampons starting on day 2 of their period, exclusively using the provided tampons (Tampax Pearl Lite). For method comparison (Figure 1), only two tampons were collected per kit. Collected tampons were fully saturated with menstrual blood before placement into transport buffer (Droplet Health). Tampons were either transported by USPS (remote), courier (local) or mobile phlebotomist (local).

#### 3.2.4 Sample Shipment and Tracking

All samples were transported at ambient temperature. US Postal Service (USPS) Priority shipping was used for all remote subjects; these samples were tracked using shipping time stamps provided by the USPS. Samples from local subjects were transported either by mobile phlebotomist or a separate courier.

#### 3.2.5 Temperature Tracking

For 35 remote donors, a USB-based temperature tracking system (TZone Digital Technology Co.) was employed.

#### 3.2.6 HbA1c Quantification

The FDA-approved and NGSP-approved Bio-Rad Variant II Turbo machine was employed for %HbA1c quantification via ion-exchange high performance liquid chromatography (HPLC). This instrument was employed both for menstrual and venous derived hemoglobin.

### 3.3 Comparison of HbA1c Measurements

Method comparison was performed by comparing tampon HbA1c versus venous blood (N=52), with samples collected on day 2 of the menstrual period. Samples were processed within 72 hours of collection. A minimum of 6 patients were recruited from each of three HbA1c ranges: <5.7, 5.7-6.4, and >6.4.

### 3.4 Stability Testing

Stability testing was performed at a range of temperature and time conditions to demonstrate robustness in practical transport conditions. A total of 12 subjects were recruited evenly across each of the following HbA1c ranges: less than 5.7, 5.7-6.4, and 6.4. Samples were processed within 18 hours of collection using local venous donors only.

Each received tampon was divided into two fragments (T0 and T1) along the long-axis seam using an ethanol-sterilize razor blade, before incubation. The HbA1c for T0 was quantified immediately. The T1 fragment was placed into a VWR Personal Low Temperature Incubator (VWR Catalog number 89511-416) with varying temperature profiles depending on condition. Exact subject counts and temperature conditions are summarized in Table 2.

**TABLE 1.** Donor types used in the method comparison and stability analysis.

| Donor | Collection | Shipment | Cooling | Samples included in analysis |
| --- | --- | --- | --- | --- |
| Local | Mobile phlebotomist | Courier | None | 1-3 tampons, 3-4 mL venous blood in one EDTA (purple top) tube |
| Remote | Self-collected | USPS | None | 1-3 tampons, 10 uL capillary blood in capillary collection tube |

USPS, United States Postal Service; EDTA, Ethylenediaminetetraacetic acid.

**TABLE 2.**
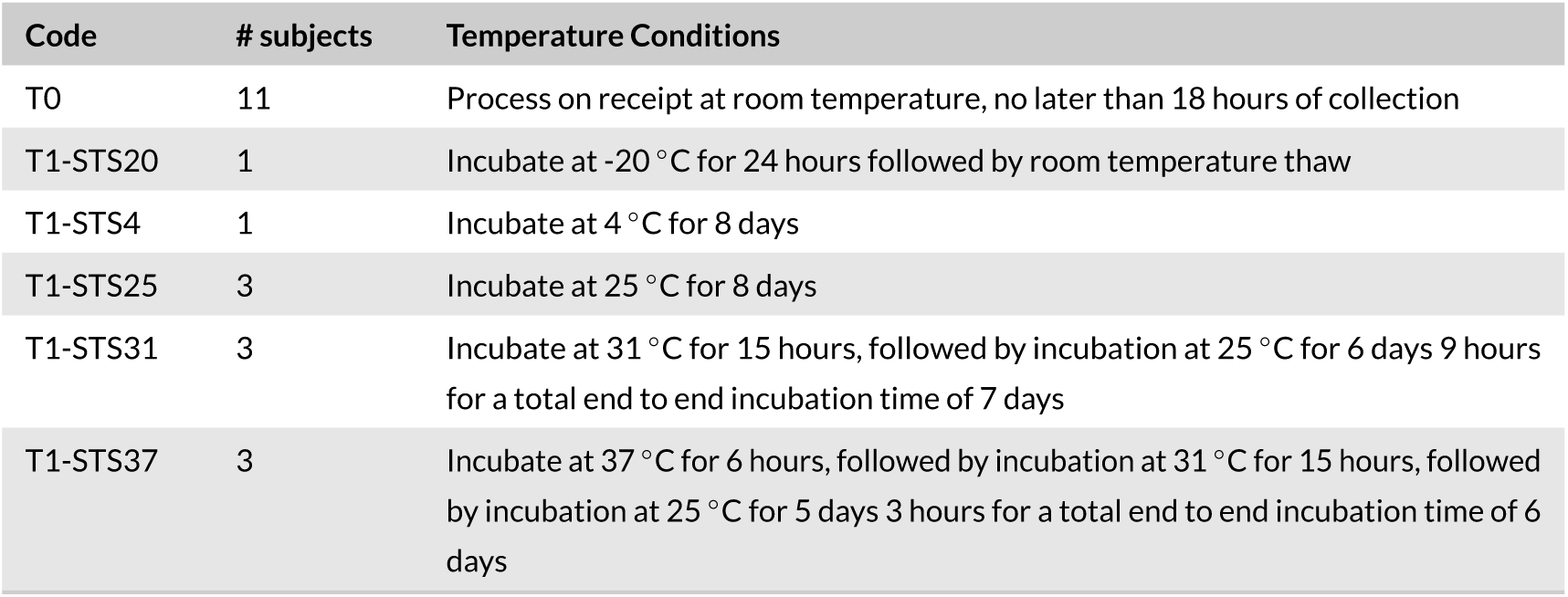
Donor types used in the method comparison and stability analysis.a

### 3.5 Interference Testing

Interference testing was performed to assess whether common exogenous and endogenous components of the vaginal and uterine environment could interfere with HbA1c assessment, including the tampon material (Tampax Pearl Lite), mucin (M3895-100MG, Sigma-Aldrich), triglycerides (glyceryl trioleate and triolein, INT-01T, Sun Diagnostics, LLC), Astroglide Liquid (water-based, over-the-counter), and Monistat-7 Complete Therapy Creme (over-the-counter). A total of N=5 subjects were employed, with samples from each subject were tested across each of five interference conditions - the tampon substance itself, and four chemical interferents. For the tampon interferent, venous blood from the corresponding patients was employed, since that blood had not experienced prior contact with tampon material. To assess tampon interference, 1 mL of venous blood was added one-quarter of a tampon in a transport container and incubated at 37 °C for 4 hours.

For other interferents, menstrual blood was used. Interferent concentration was 1000mg/dL for triglycerides and 1% w/v for mucin, Astroglide, and Monistat. Menstrual samples and chemical interferents were mixed and incubated for 1 hour at 37 °C before processing to compute menstrual HbA1c.

### 3.6 Within-Kit Tampon Agreement

To assess whether reported HbA1c values tampons were sufficiently consistent, we first plotted HbA1c values obtained from the first-collected versus the second-collected tampon. A linear regression analysis was performed using thev Regression tool under the Data Analysis panel in Excel Version 16.27 (Office 365).

Second, we compared the overall distribution of HbA1c values obtained in a worst-case scenario where the lower-valued tampon is always reported versus when the higher-valued tampon is reported from a kit. Separate empirical distributions were constructed across the first 47 kits with two available tampons, composed of only the higher-valued tampon, and separately of only the lower-valued tampon. To compare these two distributions, a 2-sample K-S test was performed.

### 3.7 Evaluation of Accuracy in the Diagnosis of Prediabetes

To evaluate whether menstrual blood performs comparably to venous blood in the diagnosis of prediabetes (Figure 2E), we calculated the true positive and false positive rates using the same 63 tampons from 63 subjects in the method comparison data, by varying the diagnostic threshold for menstrual HbA1c in determining prediabetes. We defined the optimal diagnostic threshold based on the value that minimized the sum of false positive and negative rates.

### 3.8 Experience Survey

Subjects were selected to receive a survey at random from a subset of existing study patients with HbA1c less than 6.5 as confirmed by venous or capillary blood methods. A total of 81 received the survey, and 63 subjects completed the survey. The survey contained five questions, subjects were required to submit answers to all 5 questions in order to complete the survey, and subjects who completed their survey received $5 in Amazon gift cards. The survey is as follows:

1. Which of these blood collection methods have you tried? (Check all that apply)
  a. Tampon
  b. Venous blood draw (usually in your arm)
2. If your doctor asks you to complete a blood test, which of these would you prefer? Please choose one.
  a. Tampon blood test by mail
  b. Venous blood test at your local outpatient lab
3. If your doctor asks you to do repeated blood test every three months, which one of these tests would you most likely do? Please choose one.
  a. Tampon blood test by mail
  b. Venous blood test at your local outpatient lab
4. 4. How likely are you to recommend a tampon blood test? (1 not at all likely, 10 Extremely Likely)
5. 5. How likely are you to recommend a venous blood test? (1 not at all likely, 10 Extremely Likely)

Scores for questions 4 and 5 were bucketed for analysis: detractors (1-6), passives (7-8), and promoters (9-10). Differences in proportions of detractors, passives, and promoters between tampon and venous options were examined using a two-proportion z test (Figure 4A). Differences in proportions of tampon versus venous test (Figure 4B) were examined using a one-proportion z test.

### 3.9 Geographic Distribution of Prediabetic Incidence

Projections of geographic distribution of prediabetic incidence as a percentage of the state population (Figure 3A) was computed based on the distribution of diabetes incidence based on the 2017 National Diabetes Statistics Report released by the CDC. To estimate the geographic distribution of prediabetes, we first obtained the state-by-state incidence of diabetes, the national incidence of diabetes, and the national incidence of prediabetes. We then applied this formula to each state:

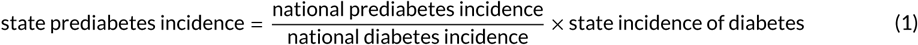

## ACKNOWLEDGMENTS

We would like to thank Rengaswamy Srinivasan PhD for discussions on chemistry, Horatio Fung Pharm D for consultation in Pharmacology, Sue Ling Fung MS for discussions on laboratory science, Cody Ebberson MBA for discussions on healthcare informatics, and Randie R. Little PhD of NGSP for detailed manuscript review and comments. JOA is funded by the Doris Duke Charitable Foundation and NYU Langone Health.

